# NeuroGate: waveform translation between brain regions

**DOI:** 10.64898/2026.09.09.750499

**Authors:** Ali Zareh, Rufeyda Yağcı, Mehmet Kemal Özdemir

**Author notes:** Contributing authors.

## Abstract

The brain processes information across distributed circuits, yet a typical experiment records only a few regions, leaving the rest unobserved. Connectivity and latent-embedding methods relate brain regions but do not return the waveform of an unrecorded one. Here we introduce NeuroGate, a framework for cross-regional neural signal translation that recovers an unrecorded region’s waveform from a recorded one. Across 56 pathways spanning rodent LFP, human sEEG and ECoG, and scalp EEG, NeuroGate predicts waveforms more accurately than 13 deep-learning and linear baselines, with low across-session variance and high median accuracy. We tested the predictions against a circuit perturbation: silencing piriform-cortex output with tetanus toxin light chain collapsed piriform-to-bulb predictions in 8 of 8 folds while the reverse was preserved, matching the monosynaptic anatomy. We demonstrate two applications: quantifying how recording redundancy bounds the information from each added source, and a reusable perturbation protocol for verifying directional translation in any circuit.

---

Behavior emerges from interactions across many connected brain regions [1, 2], yet every recording technology captures only a small fraction of these regions. Human electrode placement is dictated by clinical indications and chronically lacks coverage of several regions [3, 4], while animal multi-region recordings are restricted to experimentally targeted regions [5]. Because connected regions form circuits that exchange information through structured, bidirectional dynamics, that statistical dependence in principle allows activity in an unrecorded region to be estimated from a recorded one [6, 7]. Predicting the continuous field potential of one region from another is therefore a precondition for studying circuits that simultaneous coverage cannot reach. We frame this as *cross-regional neural signal translation* — a supervised, waveform-to-waveform mapping that recovers the local field potential (LFP), electrocorticography (ECoG), or electroencephalography (EEG) of a target region directly from a source region’s signal.

Computational tools such as LFADS [8], CEBRA [9], NDT [10], and Kraken-coder [11] transform neural activity into latent embeddings, while classical connectivity analyses [12–14] quantify pairwise interactions in frequency, power, or time domains. More recent approaches infer effective connectivity by perturbing surrogate networks [15] or predict distributed cortical activity from stimuli (TRIBE v2) [16]. All operate in the latent or low-temporal-resolution regime; none recovers the extracellular waveform itself, a precondition for any analysis on the time-domain signal. Cross-regional neural signal translation delivers a time-domain prediction of the waveform at an unrecorded region at the temporal resolution of the source recording.

We propose NeuroGate, a deep learning model trained and evaluated across rodent and human datasets spanning LFP, Stereoelectroencephalography (sEEG), ECoG, and scalp EEG under within-subject validation and leave-one-subject-out (LOSO) cross-subject testing. Three results establish the validity of this approach. First, NeuroGate generalizes across unseen subjects: the median Test−Val gap is ≈0.04 across pathways, indicating a region-to-region rather than subject-specific mapping. Second, NeuroGate predicts the target region’s waveform more accurately than 9 deep-learning and 4 linear baselines, with markedly lower across-session variance than every deep-learning baseline due to architectural design (CV = 0.22 vs. 0.47–1.55; one-sided one-sample Wilcoxon *P* = 0.002). Third, the predictions respond to circuit perturbation: when piriform cortex output is eliminated via Tetanus toxin Light Chain (TeLC) silencing [17–19], NeuroGate’s PCx→OB predictions collapse while OB→PCx is preserved, matching the monosynaptic anatomy (paired cross-subject LOSO, 8/8 folds, one-sided Wilcoxon *P* = 0.004, *d*_*z*_ = 1.62; label-permutation *P* = 0.0099).

NeuroGate also extends to many-to-one mappings: in scalp EEG, prediction quality improves with the number of source regions, consistent with biophysical predictions [20]. Once trained, NeuroGate predicts the missing region’s signal in any later session that lacks it. We then demonstrate two applications: a multi-source analysis quantifies how recording redundancy bounds the information from each added region, and the TeLC experiment doubles as a reusable perturbation protocol for verifying directional translation in any circuit. NeuroGate thus enables circuit-level analyses from existing multi-region recordings; we release an open reference implementation.

## Results

### NeuroGate translates rodent LFP with low across-session variance

We developed NeuroGate on the rodent OB→PCx pathway of the Bolding & Franks CRCNS *pcx-1* cohort [19], which provides simultaneous bilateral LFP on a sensory circuit, suitable for architecture selection. Cross-region translation on the rodent OB→PCx pathway separated the thirteen baselines we trained alongside NeuroGate into two failure modes, and Fig. 1c shows that NeuroGate avoids both. The deep-learning diamonds sit close to NeuroGate’s mean but extend long whiskers below, the signature of an architecture class whose best and worst folds disagree by a factor of two; the four linear-baseline diamonds cluster tightly but at a lower mean, plateauing near *R*^2^ ≈ 0.38. Held-out *R*^2^ for NeuroGate was 0.517 ±0.113 across nine leave-one-session-out folds of the CRCNS *pcx-1* cohort [19], well above the natural OB↔PCx coupling baseline of *R*^2^ = 0.115. We report *R*^2^ for a prediction against its target and Pearson *r* between two signals (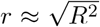 for an amplitude-matched prediction); NeuroGate’s *R*^2^ = 0.517 is *r* ≈ 0.72. The two families fail on different axes, and we tested NeuroGate against each on its failing axis. The nine deep-learning baselines fail on across-session stability: NeuroGate’s across-fold coefficient of variation (CV = 0.22) was the lowest of all ten architectures, with every deep-learning baseline’s per-architecture CV lying above it (range 0.47–1.55; one-sided one-sample Wilcoxon, *P* = 0.002). The four linear baselines are stable across sessions but fail in accuracy: NeuroGate exceeded each in all nine folds (Δ*R*^2^ ≈ +0.14 per baseline; paired one-sided Wilcoxon *P* = 0.002 each). NeuroGate is thus decisively the most stable across sessions, and more accurate than every linear baseline in every fold; the deep-learning baselines ran in their published configurations, so we rest that comparison on stability rather than mean accuracy.

**Fig. 1.**
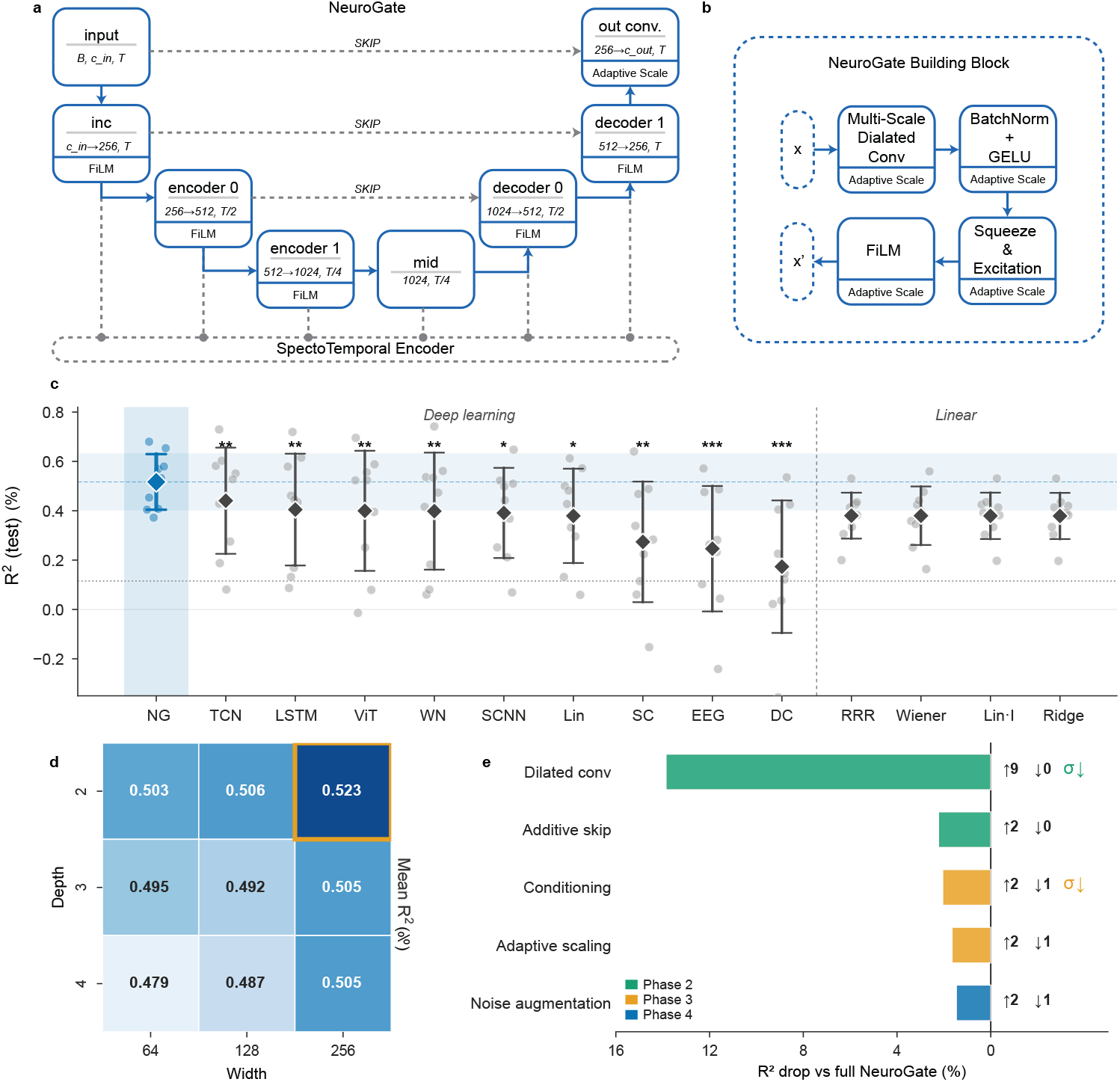
NeuroGate architecture, baseline comparison, hyper-parameter tuning, and per-component ablation. **a)** NeuroGate model overview. Encoder–decoder backbone (input inc encoder 0→encoder 1→mid→decoder 1→decoder→0→out conv) with three SKIP connections. A SpectroTemporal Encoder produces a global conditioning vector injected at every block via Feature-wise Linear Modulation (FiLM, Adaptive Scale); channel widths and temporal subsampling factors annotated on each block. **b)** NeuroGate Building Block: Multi-Scale Dilated Conv → BatchNorm + GELU → Squeeze-and-Excitation → FiLM, producing the residual update *x*^*′*^ from input *x*. **c)** Held-out *R*^2^(test) on OB→PCx (CRCNS pcx-1; *n* = 9 leave-one-session-out folds) for NeuroGate (NG, highlighted) and 13 alternatives — 9 deep-learning (TCN, temporal convolutional network [21]; LSTM [22]; ViT, 1-D vision transformer [23]; WN, WaveNet [24]; SCNN, SimpleCNN; Lin, deep linear projection; SC, ShallowConvNet [25]; EEG, EEGNet [26]; DC, DeepConv [25]) and 4 linear baselines (RRR, reduced-rank regression [27]; Wiener, frequency-domain Wiener filter; Lin-I, linear-instant least-squares; Ridge, ridge-instant regression). Grey dots, per-session *R*^2^; diamonds, mean; whiskers ±1 s.d. Pale-blue band, NG mean ±1 s.d.; dotted line, natural OB↔PCx coupling ceiling (*R*^2^ = 0.115). Asterisks: SD ratio *σ*_other_*/σ*_NG_ (∗ ≥ 1.3×, ∗∗ ≥1.7×, ∗∗∗ ≥2.2×). NeuroGate’s across-fold coefficient of variation (CV = 0.22) is the lowest of all ten architectures; every deep-learning baseline’s per-architecture CV (range 0.47–1.55) lies above it (one-sided one-sample Wilcoxon, *P* = 0.002), and NeuroGate exceeds every linear baseline on *R*^2^ in all nine folds (paired Wilcoxon *P* = 0.002). **d)** Depth × Width tuning grid (*n* = 9 sessions per cell); cell value = mean test *R*^2^. Orange border marks the carried-forward configuration (depth = 2, width = 256; *R*^2^ = 0.523). **e)** Per-component leave-one-component-out (LOCO) contribution to *R*^2^, as percent drop vs the matched-fold full-NeuroGate baseline (*n* = 9 sessions; full-NG mean *R*^2^ = 0.523). Bar colour: green Phase 2; orange Phase 3; blue Phase 4. Right of each bar, ↑*n* / = *n* / ↓*n* per-fold counts of sessions where removal improved, was neutral, or hurt the model (*τ* = 0.02 *R*^2^); *σ* ↓marks variance-stabilisers. Dilated conv dominates (Δ*R*^2^ ≈+0.07; 9/9 sessions improved). Source data: exact per-fold, per-pathway, and per-experiment values for every panel of Figs. 1–3 are provided as Supplementary Tables S1–S8; panels here draw on Supplementary Tables S2 (baseline comparison), S3 (leave-one-component-out ablation), S4 (depth×width sweep), and S7 (natural OB↔PCx coupling).

A nine-cell depth–width sweep selected depth = 2, width = 256 as the carried-forward configuration (sweep-cell mean *R*^2^ = 0.523; per-fold LOSO evaluation reports 0.517; Fig. 1d). Because this configuration was selected on the rodent OB→PCx data, the pcx-1 *R*^2^ of 0.517 is a mildly optimistic within-pathway estimate; the five further datasets of the next section use this architecture without re-tuning and supply unbiased, out-of-selection evidence. The grid is a monotone descent: shallower-and-wider configurations generalised better than deeper-and-narrower ones at every width, with *R*^2^ declining cleanly from depth = 2 to 4. Per-component leave-one-component-out ablation attributed the largest drop to the multi-scale dilated-convolution block (Δ*R*^2^ = ≈ +0.072, 9 of 9 sessions improved, paired Wilcoxon *P* = 0.004; Fig. 1e). The same block stabilized the across-session distribution (*σ* ratio 1.10 versus full NeuroGate). FiLM conditioning, additive skip connections, adaptive scaling, and noise augmentation each contributed modest performance gains (Δ*R*^2^ = 0.008–0.012). Noise augmentation, the smallest contributor (Δ*R*^2^ +0.008), is the one augmentation the deep-learning baselines omit by convention; removing it did not close the variance gap of Fig. 1c, so that gap is not an augmentation artefact. We then asked whether the same architecture carried to other modalities and pathways.

### Cross-region translation generalises across rodent, intracranial, and scalp recordings

To test whether NeuroGate converges across modalities, species, and spatial resolutions, we evaluated it on six public datasets: rodent LFP (CRCNS *pcx-1* OB↔PCx [19], *n* = 9 sessions; rodent PFC↔CA1 [28], *n* = 13), human sEEG (Boran intracranial medial temporal lobe [29], *n* = 9 patients), ECoG (Miller motor-imagery [30], *n* = 7 subjects), and two scalp-EEG cohorts (P300-oddball [31], *n* = 13; CMx7-EEG motor-imagery [32], *n* = 12). Every model was evaluated under cross-subject leave-one-subject-out (Test) and within-subject (Val) regimes, each pathway compared by one-sided paired Wilcoxon signed-rank test against the natural inter-region coupling baseline — a model-free reference for how much of the target a source explains through raw shared variance, distinct from the thirteen trained baselines of Fig. 1c. Across the 56 directed pathways, NeuroGate’s predictions exceeded the natural-coupling baseline on 45 pathways under Test and 55 under Val, with a median test *r* of 0.53 (range 0.26–0.82; Fig. 2a).

**Fig. 2.**
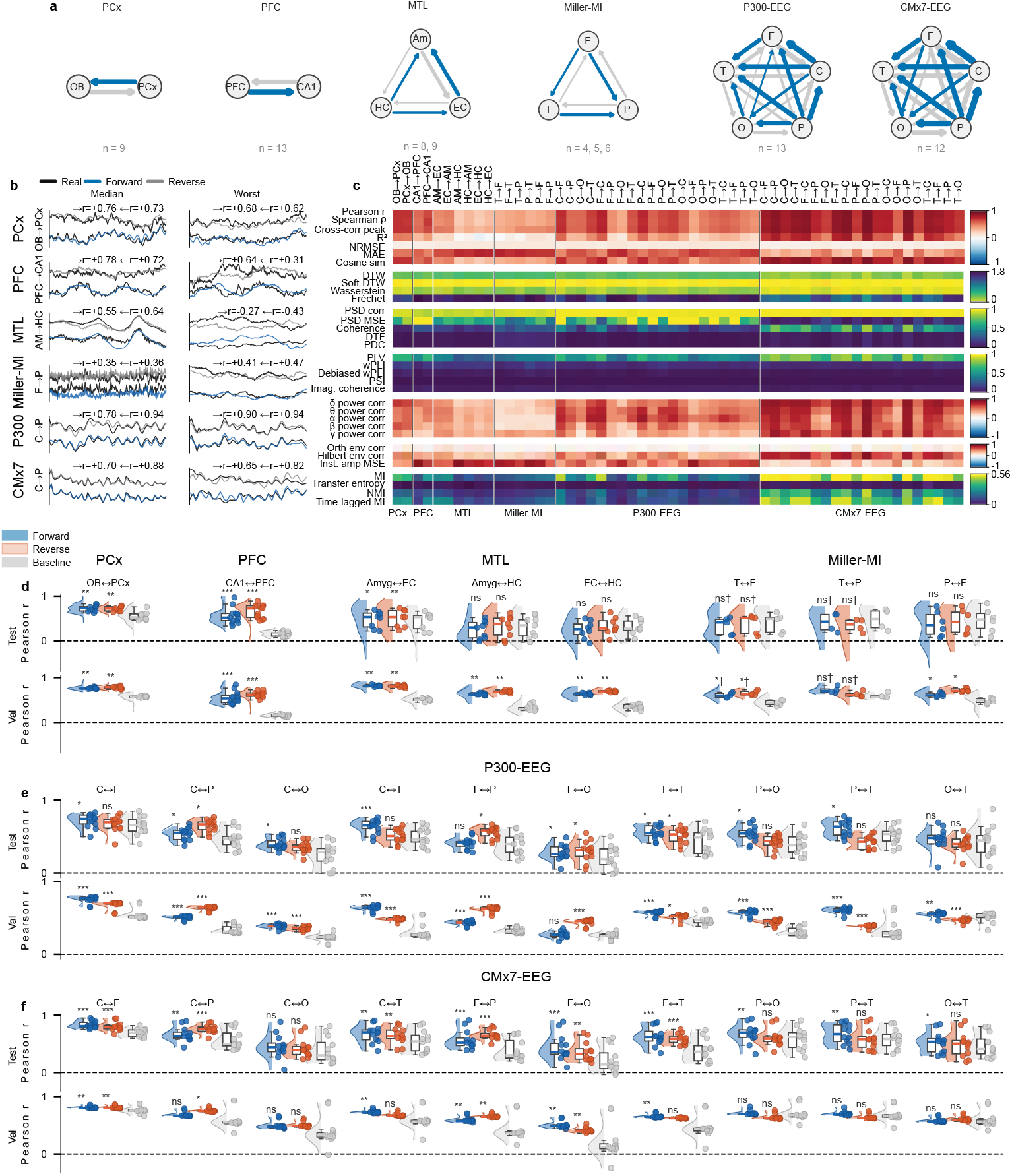
NeuroGate cross-region Pearson *r* across six datasets — networks, traces, multi-metric fidelity, and per-pathway distributions. **a)** Six directional networks (P300-EEG, CMx7-EEG, MTL, PCx, PFC, Miller-MI); per pair the stronger direction is blue, weaker grey, arrow thickness ∝ *r*. Per-dataset *n* below. **b)** Median (“Typical”) and worst-subject real vs predicted exemplars across six dataset rows; 250 ms *z*-scored windows with per-cell forward / reverse *r*. **c)** 33-metric reconstruction-fidelity heatmap split by dataset; seven family-specific colour scales. Every family — including phase coupling — is scored here purely as a reconstruction-fidelity metric, comparing a prediction against its own target, not as a downstream phase or oscillation analysis of the predicted signal. Hatched grey = metric not computable for that pathway. **d**–**f)** Per-pathway Pearson *r* rain-clouds (Forward blue, Reverse vermilion, Baseline grey = real-vs-real natural inter-region coupling); Test top, Val bottom. Asterisks: per-pathway one-sided Wilcoxon *r*_model_ *> r*_baseline_ (∗ *P <* 0.05, ∗∗*P <* 0.01, ∗∗∗*P <* 0.001, ns otherwise; † = *n <* 6); each pathway tested independently. **d)** rodent / sEEG / ECoG; **e)** P300-EEG; **f)** CMx7-EEG. *n* per pathway and dataset citations in Methods. Source data: Supplementary Table S1 (per-pathway reconstruction statistics — 112 pathway × direction × split cells).

The directional networks of Fig. 2a summarise per-pathway prediction strength: the strongest are the centrally anchored scalp pathways and the rodent OB↔PCx and PFC↔CA1 axes, with lighter but resolved bundles on the sparser MTL and Miller-MI grids. The waveform exemplars of Fig. 2b confirm this at the trace level: on the median held-out subject the predictions track the dominant rhythm and the timing of prominent transients, and on the worst-*r* subject the envelope is preserved while fast transients soften. Forward and reverse predictions stay visibly distinct within every cell, so the model is not collapsing the two directions onto a shared latent trajectory.

A supervised translator earns its place only if it recovers structure beyond the raw shared variance between two regions; the raincloud panels of Fig. 2d–f make this visible, pairing every forward and reverse prediction cloud with a third *Baseline* cloud — the real-vs-real inter-region coupling on the same subjects, channels, and windows used to score the model. On most pair-directions the forward cloud sits above baseline, and panel-level Wilcoxon tests on per-pathway means confirm the separation: *P <* 0.01 on the central → frontal direction in both scalp cohorts (Fig. 2e–f), *P* ≤ 0.0039 per direction × split on rodent OB↔PCx (*n* = 9; Fig. 2d), and *P <* 0.001 on rodent PFC CA1 in both directions and both splits (*n* = 13). The combined rodent + intracranial panel (Fig. 2d) is the most heterogeneous: the rodent OB↔PCx and PFC↔CA1 axes carry forward predictions well above baseline, whereas on the sparser intracranial contacts forward and baseline overlap and the contrast does not always reach significance. The Test and Val subrows remain near-identical across every panel, ruling out a test-split artefact. Across panels, NeuroGate closes the gap between what natural coupling already explains and an empirical ceiling on any input-bounded reconstruction, rather than beating natural coupling by a fixed margin on every pair.

The median Test–Val gap across the 56 pathways was ≈0.04. We read this as a positive feature: validation rewards memorising within-subject structure while test, evaluating cross-subject transfer, does not, so a small gap points to subject-invariant rather than subject-specific learning. Reconstruction so far rested on Pearson *r*; we asked whether the picture survives other similarity definitions. The 33-metric heatmap of Fig. 2c re-scores all 56 pathways across seven metric families — time-domain fidelity, shape distance, spectral amplitude, phase coupling, band-limited power, envelope similarity, and information-theoretic measures. The families agree: pathways bright in the similarity, spectral, and phase-coupling rows are bright in the information-theoretic rows and darkest in the distance rows. The cross-family agreement, with the directional asymmetry of Fig. 2a, indicates that NeuroGate’s predictions carry the direction- and circuit-specific structure of each anatomy, not a summary that happens to look right under one metric. The pathway-level correlations describe how well the predictions match, not where in the source signal the predictable variance comes from.

### Application: quantifying multi-region recording redundancy

We next applied NeuroGate as an experimental-design tool, asking how many regions a study must record to recover a given target. In both scalp cohorts, the target could be reconstructed from any one of four neighbouring regions, with each additional source adding less than the last (Fig. 3a, 3d). The saturation curves rise sharply from *k* = 1 to *k* = 2, bend at *k* = 3, and flatten by *k* = 4, the same shape across both cohorts. Per-target *R*^2^(*k*) for *k* ∈ {1 … 4} sources fit the geometric form 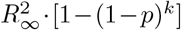, with *p* as a phenomenological summary of saturation-curve shape rather than a measure of statistical independence — a two-parameter fit to four points (per-target values in Supplementary Table S8). Mean *p v* 0.59 in P300-EEG (*n* = 13) [31] and ≈0.63 in CMx7-EEG (*n* = 12) [32], giving 1 − *p* ≈ 0.40 as a curve-shape descriptor. Roughly four-tenths of each new source’s marginal contribution overlapped with what earlier sources already supplied, consistent with the saturation profile expected for scalp recordings sharing common deep generators [20].

**Fig. 3.**
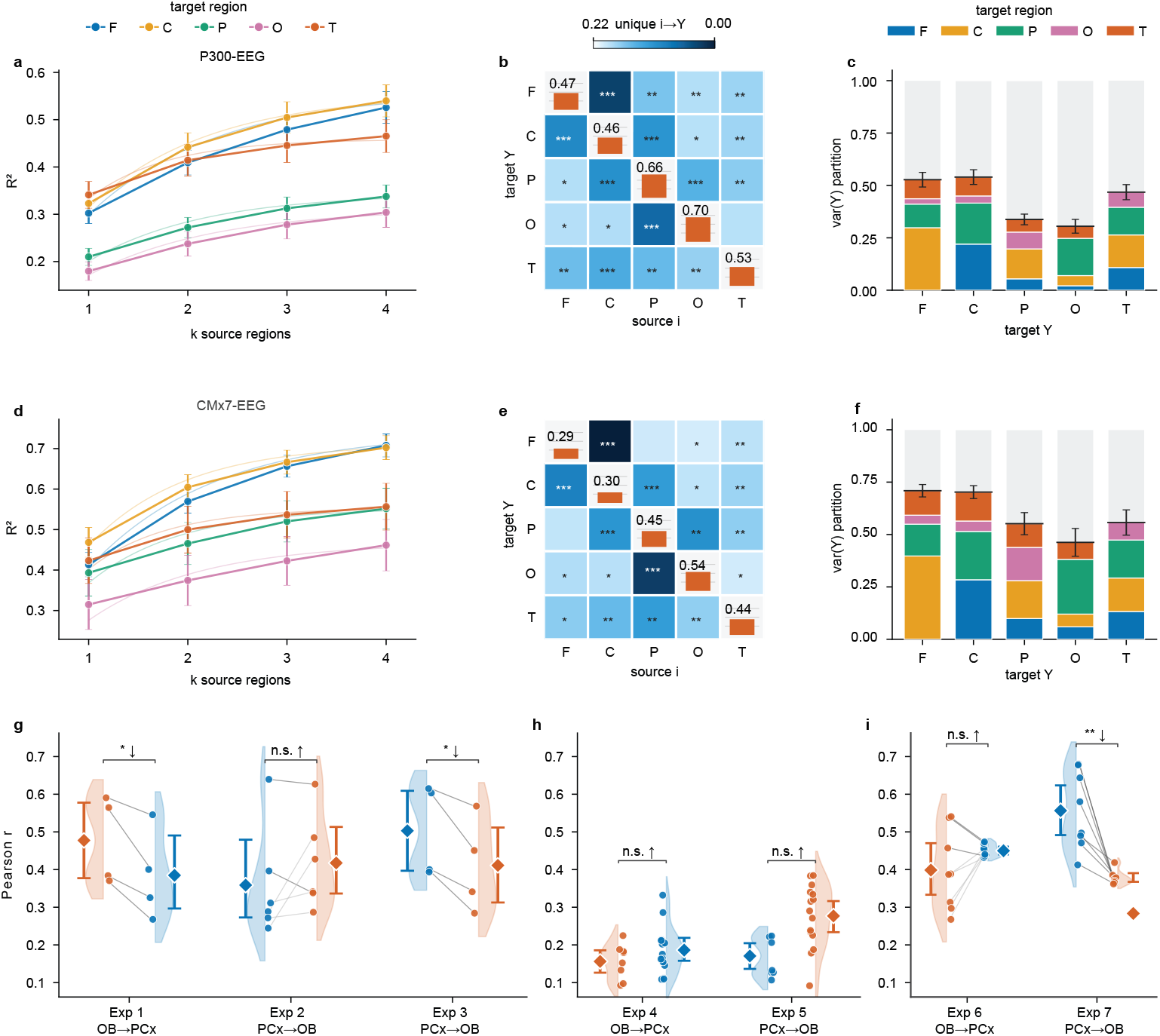
Two applications of NeuroGate: a multi-region recording-redundancy analysis and a perturbation protocol that recovers directional anatomy. Significance throughout: ∗ *P <* 0.05, ∗∗ *P <* 0.01, ∗∗∗ *P <* 0.001, ns otherwise; †= *n <* 6. Top two rows (**a**–**f**) decompose predictions into IT atoms on P300-EEG (**a**–**c**, *n* = 13) and CMx7-EEG (**d**–**f**, *n* = 12) across five scalp regions (F frontal, C central, P parietal, O occipital, T temporal); raw-waveform multi-source predictions from end-to-end NeuroGate. **a), d)** *R*^2^(*k*) saturation ladder for *k* ∈ {1 … 4} active sources; markers, mean ± s.e.m.; light dashed lines, per-target geometric fit 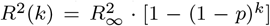. Per-source *p* = 0.59 P300 / 0.63 CMx7 → redundancy *R* ≈ 0.40. **b), e)** 5 × 5 source × target unique-contribution matrix (rows = target *Y*, columns = source *i*). Off-diagonal: leave-one-out unique *a*_*i*→*Y*_ ; shared power-law colourbar (*γ* = 0.55). Per-cell one-sample Wilcoxon *a*_*i*→*Y*_ *>* 0 with BH-FDR within panel. Diagonal: intrinsic 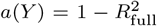 as vermilion bar (0–1) over neutral grey; numeric value above. **c), f)** Per-target IT decomposition stacked bar: four Shapley atoms (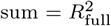; black tick = top of coloured stack) plus a grey *a*(*Y*) topper. Whisker, ± s.e.m. of 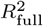. Bottom row (**g**–**i**) — rodent OB↔PCx under TeLC silencing. Blue = Ctrl, vermilion = TeLC; per-group raincloud; bracket: per-pathway one-sided Wilcoxon *r*_model_ *> r*_baseline_. **g)** TeLC unilateral-injection cohort, simul-trained: Exp 1 OB→PCx (*n* = 4, paired-*t P* = 0.020, Wilcoxon *P* = 0.063, *d*_*z*_ = 1.72); Exp 2 PCx→OB (*n* = 6, paired Wilcoxon, ns); Exp 3 Y-junction (*n* = 4, paired-*t P* = 0.016, Wilcoxon *P* = 0.063, *d*_*z*_ = 1.89). **h)** Thy1-ChR2 × Emx1-Cre TeLC cohort, simul-trained, between-subjects (8 vs 15 recordings): Exp 4, Exp 5 both ns. **i)** Same line, NeuroGate fine-tuned per fold (LOSO): Exp 6 OB→PCx (*n* = 8, ns); Exp 7 PCx→OB headline (**8/8 direction-consistent**, paired Wilcoxon *P* = 0.004, *d*_*z*_ = 1.62, percentile bootstrap 95% CI [1.22, 3.02], 10,000 resamples, seed 42; label-permutation *P* = 0.0099, *N* = 100, observed *k* = 8 vs null *k* ∈ [0, 7] — Supplementary Table S6). Source data: Supplementary Tables S5 (TeLC experiments Exp 1–7) and S8 (multi-source decomposition); the Exp 7 label-permutation null is Supplementary Table S6.

We localised source-by-source contributions through the 5 × 5 unique-contribution matrix per dataset (Fig. 3b, 3e). Off-diagonal cells report each source’s leave-one-out unique contribution *a*_*i*→*Y*_ after the other three sources are accounted for; diagonal cells report the intrinsic share 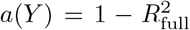, which measures the variance in *Y* that the full four-source model fails to explain. The matrix is dominated by mid-range off-diagonal cells in both cohorts, brighter along the central-region rows; most off-diagonal cells passed BH-FDR within panel (*q <* 0.05), so each source carried unique information beyond what the other three provided. Per-target Shapley decomposition (Fig. 3c, 3f) splits the predictable variance into ordering-invariant atoms summing to *R*^2^, with the intrinsic share added as a grey topper. The bars divide unevenly: central targets carry slim grey toppers above four comparable atoms, peripheral targets tall toppers above smaller atoms (central *a*(*Y*) ≈ 0.45 versus occipital ≈ 0.70 in P300-EEG; ≈ 0.31 versus ≈ 0.53 in CMx7-EEG).

The asymptotic ceiling 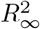 varied across targets between roughly 0.1 and 0.9, so some scalp regions are intrinsically more recoverable than others even with full input. The decomposition shows where the predictable variance comes from, not whether the predictions track directional circuit communication – a question for a perturbation.

### Application: a perturbation protocol that recovers directional anatomy

As a second application, we used NeuroGate as a perturbation protocol that isolates one directed edge at a time and, therefore, applies to a circuit of any size; we validated it here on a single causally silenced pathway, testing whether the predicted waveforms recover its known directional anatomy. Silencing piriform-cortex output with tetanus toxin light chain collapsed PCx→OB predictions in 8 of 8 held-out recordings while leaving OB→PCx intact (Fig. 3i). The seven experiments span two TeLC-silencing cohorts and three training regimes on the Bolding & Franks 2018 dataset [17–19], each cohort silencing one piriform hemisphere unilaterally with the contralateral hemisphere as within-animal control: a TeLC unilateral-injection cohort (Exp 1–3, *n* = 4–6 per pathway) and a Thy1-ChR2 × Emx1-Cre cohort (Exp 4–7; 8 control versus 15 silenced recordings), trained pooled, between-subjects, or fine-tuned per leave-one-subject-out fold. The directional prediction was anatomy-locked: TeLC eliminates synaptic release from PCx output, so only the pathway whose source must traverse PCx output should degrade, not the reverse.

The headline result emerged under per-fold LOSO fine-tuning on the *Thy1* line: PCx→OB predictions collapsed direction-consistently in 8 of 8 held-out recordings, every Ctrl→TeLC slope in Fig. 3i tilting downward with no fold reversing (Exp 7, one-sided paired Wilcoxon *P* = 0.004, *d*_*z*_ = 1.62, percentile bootstrap 95% CI [1.22, 3.02], 10,000 resamples). On the same recordings and regime, OB→PCx predictions did not show collapse, Ctrl and TeLC clouds overlapping (Exp 6, *n* = 8, *P* = 0.93). The asymmetry between Exp 6 and Exp 7 is the within-paradigm control: a generic fine-tuning artefact would tilt both directions, TeLC silencing only one, and the data match the latter. A label-permutation control over 100 random Ctrl/TeLC assignments produced direction-consistency counts *k* centred at 3.28 (range 0–7), with no permutation reaching the observed *k* = 8*/*8 (permutation *P* = 0.0099; Supplementary Table S6).

The independent TeLC unilateral-injection cohort was mixed. The Y-junction analysis, which sources PCx→OB from the intact contralateral hemisphere, degraded in all four recordings tested (Exp 3, *n* = 4, paired-*t P* = 0.016, *d*_*z*_ = 1.89) — the same pathway and direction as Exp 7. Exp 1 and Exp 3 score the same four recordings and are not independent tests. OB→PCx also degraded (Exp 1, paired-*t P* = 0.020, Wilcoxon *P* = 0.063, *d*_*z*_ = 1.72), which runs against the monosynaptic prediction, and the direct PCx→OB comparison did not change (Exp 2, *n* = 6, *P* = 0.89). At *n* = 4 the Wilcoxon is at its 1*/*16 = 0.0625 floor; the directional claim therefore rests on Exp 7’s larger sample, with Exp 3 as same-direction support.

Under between-subjects training the effect reversed: TeLC recordings scored *above* controls (Exp 4, Δ*r* = +0.043; Exp 5, Δ*r* = +0.106; 8 versus 15 recordings). The directional effect appears only under per-fold fine-tuning. We read this as a methodological boundary: between-subjects training conflates the silencing signal with cohort-level variance, whereas per-fold fine-tuning adapts to each held-out recording’s geometry (Methods). Exp 7, with the Exp 6 negative control and the convergent Exp 1 and Exp 3 evidence in the independent cohort, establishes that NeuroGate’s directional predictions track monosynaptic information flow rather than statistical co-activation [19].

## Discussion

Recovering the waveform itself has two consequences for what the method is good for. The first is evidential: a waveform prediction can be subjected to a circuit perturbation and tested causally. The directionally selective collapse under TeLC silencing places these predictions in a different class of evidence than within-distribution generalisation. In the *Thy1* cohort, under identical LOSO fine-tuning, the PCx→OB direction collapsed in 8 of 8 folds while OB→PCx was preserved (Exp 6 vs Exp 7). The asymmetry is direction-specific rather than global, but the design does not separate channel loss from the silenced signal being out of distribution for control-tuned weights (Methods). TeLC eliminates piriform cortex output [17, 18], so only the pathway whose source must traverse PCx output fails. This anatomy-matched directionality reproduces the monosynaptic OB→PCx wiring [19]: feedforward input to PCx remains, while PCx-to-OB feedback is removed. Latent-embedding methods [8–11] cannot be tested this way: their predictions live in a learned embedding, not the waveform space of the perturbation’s effect. Recovering the waveform is therefore not a stylistic preference but a precondition for perturbation-based validation.

The second consequence is methodological, demonstrated through two applications. Because the prediction shares the waveform space of a circuit perturbation, the TeLC experiment doubles as a reusable protocol, not a one-off validation of this circuit: any laboratory can silence one node and test which direction of prediction collapses, verifying directional translation in its own pathway (Fig. 3g–i). The multi-source decomposition (Fig. 3a–f) is the second application, for experimental design: redundancy among scalp regions plateaus the marginal information in each additional source (*R* ≈ 0.40), bounding the benefit of the third or fourth source, and the largest-contributing location is identifiable from the off-diagonal cells of the leave-one-out matrix. Both applications require an electrode-grade trace at the unrecorded site. Latent-embedding pipelines [8–11] return embeddings, not waveforms; effective-connectivity inference [15] returns directed coupling estimates, not signals. Neither substitutes for the waveform itself.

NeuroGate also differs fundamentally from cross-subject imputation. SuperEEG [33] infers activity at unrecorded locations from a linear Gaussian-process estimator built on between-electrode correlation matrices pooled across people; because those matrices are symmetric, it is an undirected spatial interpolator, evaluated by how closely reconstructed traces correlate with held-out ones. NeuroGate instead learns a nonlinear, directed mapping from a specific source region’s waveform to a specific target’s, and is tested not only by reconstruction accuracy but by a circuit perturbation that abolishes one direction of that mapping — evidence a symmetric correlational model has no directional structure to be subjected to. The two are complementary: spatial smoothing fills gaps within a shared anatomical frame, while directed translation recovers what one region’s activity implies about another.

Several boundary conditions remain. At *n* = 4 (Exp 1, Exp 3) the Wilcoxon is at its 1*/*2^4^ = 0.0625 floor, so the paired *t*-test is the primary statistic; the load-bearing claim rests on the 8-fold Exp 7 LOSO and its label-permutation control (*P* = 0.0099; *k* = 8*/*8 beyond the empirical null *k* ∈ [0, 7]). Cohort coverage is uneven: rodent LFP and scalp EEG dominate, the rodent prefrontal–hippocampal [28] and Miller motor-imagery ECoG [30] datasets are smaller, and human subcortical pathways are limited to the 9-patient Boran sEEG cohort [29]. The causal validation itself is one pathway, one direction, one species; perturbation cohorts in other circuits are needed to test how broadly the directional sensitivity extrapolates. NeuroGate was trained and evaluated within recording paradigm, so transfer of a mapping across behavioural or arousal states is untested. The causal LOSO results rest on cohorts of at least eight subjects or sessions, the smallest size at which per-fold estimates were stable; smaller cohorts were not characterised. Among linear comparators we evaluated a frequency-domain Wiener filter but not multivariate autoregressive or state-space predictors, which could narrow the linear-baseline gap on pathways with strong low-order temporal structure. Finally, the geometric saturation fit (Fig. 3a, 3d) is a descriptor of curve shape fitted to four points per target, not an information-theoretic decomposition; per-source attribution is given by the Shapley atoms (Fig. 3c, 3f), and the band-limited mutual-information ceiling (Methods) is a lower bound on the full-waveform ceiling.

What changes is the treatment of unrecorded regions. Once trained on a multi-region dataset, cross-regional waveform translation becomes a routine analysis step recovering signals from circuits never simultaneously recorded; an open reference implementation (github.com/AliZareh-CoE/NeuroGate) and the protocols collected here lower the barrier to adoption.

## Methods

### Datasets

We trained and evaluated NeuroGate on six public datasets. CRCNS *pcx-1* [19]: simultaneous olfactory-bulb and piriform-cortex LFP from 9 awake-mouse sessions. Rodent PFC↔CA1 [28]: paired LFP from 13 sessions. Boran intracranial medial temporal lobe [29]: sEEG of amygdala, entorhinal cortex, and hippocampus in 9 patients during a verbal working memory task. Miller motor-imagery ECoG [30]: 7 subjects with frontal, parietal, and temporal coverage. P300-EEG [31]: 13 subjects, 64-channel Biosemi mapped onto five anatomical region groups (frontal, central, parietal, occipital, temporal). CMx7-EEG [32]: 12 subjects, same five-region mapping; only the EEG channels of the multimodal EEG/fNIRS recording were used. TeLC silencing experiments used the Thy1-ChR2 × Emx1-Cre TeLC cohort from the same Bolding & Franks dataset [19] (23 recordings; 8 control, 15 TeLC; unilateral TeLC silencing of one piriform hemisphere with the contralateral hemisphere as within-animal control) and a separate TeLC unilateral-injection cohort (*n* = 4–6 per pathway). The per-dataset session and subject counts given here are the complete set entering analysis; no further recordings or analysis windows were excluded.

### Preprocessing

All datasets passed through a single preprocessing pipeline. Recordings sampled above the canonical target rate (1 kHz for LFP and intracranial; 256 Hz for scalp EEG) were anti-alias filtered and polyphase-resampled to the target; datasets already at or below the target were used at their native rate (P300-EEG at 256 Hz; CMx7-EEG at 250 Hz). Signals were partitioned into overlapping windows and per-channel *z*-scored from each window’s own statistics at batch time, with no global normalisation across the dataset, preventing train–test leakage. Window length and stride per dataset and the scalp 10–20 montage configuration were applied via MNE-Python.

### NeuroGate architecture

NeuroGate is an encoder–decoder convolutional network [34] with multi-scale dilated convolutions [35], squeeze-and-excitation gating [36], FiLM-gated conditioning [37], and adaptive output scaling. The carried-forward configuration was depth = 2, width = 256, selected by a 9-cell depth × width sweep that maximised held-out *R*^2^ on the rodent OB→PCx pathway (*R*^2^ = 0.523). Channel widths, downsampling factors, and block-internal layout are annotated in Fig. 1a,b.

Each convolutional block applied a multi-scale dilated convolution with kernel size 7 and four parallel branches at dilations (1, 4, 16, 32), corresponding to receptive fields of 7, 25, 97, and 193 ms spanning the *γ, β, θ/α*, and *δ* frequency bands of the LFP. Branch outputs were fused by softmax-weighted averaging, gated by squeeze-and-excitation channel attention, then modulated by a FiLM gate driven by a SpectroTemporalEncoder conditioning embedding derived from the input itself. The SpectroTemporalEncoder builds this 128-d embedding from two parallel branches: a spectral branch summarising log-power over 10 frequency bands up to 100 Hz, and a temporal branch of three dilated convolutions (dilations 1, 4, 16; kernel 7), each branch projected to 64 dimensions and concatenated. The FiLM gate modulates a block feature map *x* as *x* + *σ*(*g*(*e*)) ⊙ out softmax_*C*_[*q*(*x*) ⊙ *k*(*e*) *C*^−1*/*2^] ⊙ *v*(*e*), where *e* is the conditioning embedding, *q, k, v* are learned projections. The softmax runs over the *C* channels, and *σ*(*g*(*e*)) is a learned per-channel sigmoid gate. The output head produced a residual delta on the per-channel-normalised input. A SessionAdaptiveScaling module then applied a per-channel affine correction — a scale and bias predicted by a small multilayer perceptron from each session’s per-channel input mean and standard deviation, initialised to the identity transform — which adapts the normalised prediction to session-to-session amplitude differences but does not restore absolute physical units, as both source and target are per-channel *z*-scored at training and evaluation time.

### Training

Training used AdamW (learning rate 1 × 10^−3^) with a 5-epoch linear warmup followed by cosine decay over the remaining 75 epochs (80 total). Batch size was 64; the loss was per-channel *L*_1_ on *z*-scored target windows; gradient norms were clipped at 5.0; early stopping triggered after 15 consecutive epochs without validation improvement. Stochastic noise augmentation (Gaussian, 1/*f* pink, channel dropout, temporal-bin dropout) was applied with 50% probability per batch. All random number generators (NumPy and PyTorch, CPU and CUDA) were seeded at 42; under distributed training the DataLoader sampler shuffling was reseeded each epoch. cuDNN auto-tuning was enabled for throughput during the architecture-screening sweep only, and disabled for the final NeuroGate and baseline runs, which are therefore reproducible from the seed. Architecture-screening sweeps used 8 NVIDIA A100 GPUs via PyTorch DistributedDataParallel; final NeuroGate training used a single A100 or 8×A100 with FullyShardedDataParallel. Implementations used PyTorch ≥ 2.0, NumPy ≥ 1.24, SciPy v1.11, statsmodels v0.14, scikit-learn ≥ 1.3, MNE-Python v1.6, and Python ≥ 3.10.

### Cross-validation and evaluation

Cross-validation was leave-one-session-out for the rodent datasets and leave-one-subject-out for the human datasets. Within each fold the held-out session or subject was the Test split; the remaining sessions or subjects were partitioned into Train (≈ 70%) and Val (≈ 30%) — a contiguous temporal split for the continuous rodent recordings and a subject-level holdout for the human multi-source loop — deterministic from a per-fold seed and reused across NeuroGate and the baselines for exact comparability. Multi-source raw-waveform experiments (P300-EEG and CMx7-EEG; see *Multi-source information decomposition*) used an analogous per-subject LOSO loop.

Two metrics were computed on each held-out window pool: pooled Pearson correlation *r* = cov(pred, target)*/*[std(pred) · std(target)] and pooled *R*^2^ = 1 −SS_res_*/*SS_tot_, both pooled across all output channels and time within the held-out subject or session. The natural inter-region coupling baseline reported alongside model performance was the same metric computed on the real source against the real target signal from the same simultaneous windows.

### Baseline models

We compared NeuroGate to 13 alternatives on the rodent OB→PCx pathway: nine deep-learning architectures — a temporal convolutional network [21], an LSTM [22], a 1-D vision transformer [23], a WaveNet [24], a SimpleCNN (in-house, 4-layer residual CNN), a ShallowConvNet [25], an EEGNet [26], a DeepConvNet [25], and a deep linear projection — and four linear baselines: linear-instant least-squares, ridge-instant regression with a validation-tuned *ℓ*_2_ penalty, a frequency-domain Wiener filter, and reduced-rank regression. Each deep-learning baseline was implemented at NeuroGate’s input/output channel count and trained under matched hyperparameters without noise augmentation, to match each baseline’s published reference implementation; the within-NeuroGate LOCO ablation (Fig. 1e; Results §1) confirms that augmentation contributes only Δ*R*^2^ ≈ +0.008. Each baseline therefore ran in its published reference configuration rather than under a per-baseline hyperparameter search, whereas NeuroGate’s depth and width were set by the architecture sweep. This asymmetry is real and we do not minimise it; it cannot, however, account for the variance result of Fig. 1c, because the sweep selected the configuration with the highest mean held-out *R*^2^ — not the lowest across-session variance. Mean accuracy and across-session variance are distinct objectives, and no grid cell was chosen to minimise variance, so the per-architecture variance gap cannot have been manufactured by the tuning step. Linear baselines were fit on the same Train pool, validated on Val, and evaluated on Test as NeuroGate.

### Multi-source information decomposition

On P300-EEG (*n* = 13) and CMx7-EEG (*n* = 12) we decomposed NeuroGate’s predictions into source-specific information atoms by evaluating all 15 non-empty subsets of the four scalp regions other than the target. For each subset of *k* sources, an end-to-end NeuroGate was trained with the *k* raw waveforms concatenated along the channel dimension as the input and the target’s waveform as the prediction target, under the per-subject LOSO protocol; per-(subject, target) saturation curves 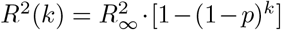 were fit, yielding per-source independence *p* and asymptotic ceiling 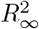.

Leave-one-out unique contributions *a*_*i*→*Y*_ = *R*^2^(others) − *R*^2^(others \ {*i*}) and the diagonal intrinsic share 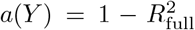 were computed per (source, target). A Shapley four-atom decomposition [38] averaged marginal contributions across all 4! source orderings; atoms summed to 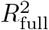 by construction. The empirical ceiling 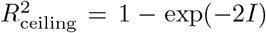 was computed from the mutual information *I* between the target and the full source set, estimated with the Kraskov–Stögbauer–Grassberger *k*-nearest-neighbour estimator (*k* = 5) [39].

### TeLC perturbation experiments

TeLC silencing data were drawn from the Bolding & Franks 2018 dataset [19] as two cohorts, both using unilateral TeLC silencing of one piriform hemisphere with the contralateral hemisphere as within-animal control: a Thy1-ChR2 × Emx1-Cre cohort (23 recordings; 8 control, 15 TeLC) and a separate TeLC unilateral-injection cohort (*n* = 4–6 per pathway). Three training regimes were applied across seven experiments. **Simul-trained:** the base rodent OB↔PCx NeuroGate was applied without modification (Exp 1, OB→PCx, *n* = 4; Exp 2, PCx→OB direct, *n* = 6; Exp 3, Y-junction, *n* = 4). **Between-subjects:** separate models were trained on Ctrl-only and TeLC-only pools (Exp 4 OB→PCx; Exp 5→PCx OB; 8 vs 15 recordings). **Per-fold LOSO fine-tuning:** for each held-out control session a copy of the base model was fine-tuned on the other 7 control sessions for 10 epochs at a learning rate of 1 × 10^−4^, then evaluated on the held-out control and on each of the 15 TeLC sessions (Exp 6, OB→PCx, *n* = 8; Exp 7, PCx→OB, *n* = 8 — the headline causal experiment). Direction-consistency was the count of folds where *r*_TeLC_ *< r*_Ctrl_. The unit of pairing in Exp 6 and Exp 7 is therefore the fold, not the animal: each fold contributes one held-out control recording and one score aggregated over the same 15 TeLC recordings, so the eight comparisons share their TeLC arm and are not mutually independent. The label-permutation control of Exp 7 was run through the identical pipeline and does not assume independence across folds. Because fine-tuning is performed on control sessions only, the held-out control is in-domain for the fine-tuned weights while the TeLC recordings are not; the measured gap therefore includes any differential benefit of fine-tuning as well as any effect of silencing, and the two are not separated by this design.

The Y-junction analysis (Exp 3) re-routes the PCx→OB pathway through the contralateral, non-injected piriform hemisphere as the source, separating the source-side recording from the silenced output pathway. A label-permutation shuffle control on Exp 7 (100 random Ctrl/TeLC assignments through the same fine-tuning pipeline; master seed 42) returned permutation *P* = 0.0099 with the observed *k* = 8*/*8 direction-consistency beyond the maximum of the empirical null distribution (*k*_null_ ∈ [0, 7]); see Supplementary Table S6.

### Statistics

All tests are reported as (test name, n, exact *P*, effect size, software function). Per-pathway model-vs-baseline comparisons (Fig. 2d–f): one-sided paired Wilcoxon signed-rank on per-subject (model *r* − baseline *r*) on held-out windows (`scipy.stats.wilcoxon(alternative=`‘`greater`’`, zero method=`‘`wilcox`’)); no FDR across pathways because each pair is an independent scientific question. NeuroGate vs the deep-learning baselines on across-session stability (Fig. 1c): the across-fold coefficient of variation (CV = SD / mean of held-out *R*^2^) was computed separately for NeuroGate and for each of the nine deep-learning architectures, and NeuroGate’s CV was compared against the nine per-architecture baseline CVs by a one-sided one-sample Wilcoxon signed-rank test (*n* = 9 architectures). NeuroGate vs the four linear baselines on accuracy: per-fold held-out *R*^2^ was compared by a one-sided paired Wilcoxon signed-rank test across the nine leave-one-session-out folds (*n* = 9 folds). Per-component LOCO ablation (Fig. 1e): paired Wilcoxon per component; the dilated-conv claim rests on effect size and per-fold sign consistency rather than the raw *P*.

#### Paired *n* = 4 cohorts (Exp 1 and Exp 3)

a paired one-sided *t*-test (`scipy.stats.ttest rel(alternative=`‘`greater`’)) as the primary statistic, with the nonparametric Wilcoxon reported alongside for transparency. Exp 6 and Exp 7 paired LOSO (*n* = 8): one-sided paired Wilcoxon. Exp 4 and Exp 5 between-subjects: one-sided Mann–Whitney *U* (`scipy.stats.mannwhitneyu(alternative=`‘`greater`’)). The Exp 7 label-permutation shuffle control: master seed 42, *N* = 100 random Ctrl/TeLC label assignments through the same fine-tuning pipeline; permutation *P* = (#{*k*_perm_ ≥ *k*_obs_} + 1)*/*(*N* + 1) per Phipson & Smyth 2010. Multi-source unique-contribution off-diagonals (Fig. 3b, 3e): one-sample Wilcoxon *a*_*i*→*Y*_ *>* 0 with Benjamini–Hochberg FDR control within panel at *q <* 0.05 (`statsmodels.stats.multitest.multipletests(method=`‘`fdr bh`’)) [40]. Bootstrap confidence intervals [41] used 10,000 resamples with the percentile method.

## Data availability

Datasets: CRCNS *pcx-1* [19], G-Node `10.12751/g-node.d76994` [29], OpenNeuro `ds007554` [32], the Miller motor-imagery ECoG library [30], and OpenNeuro `ds003061` [31] are publicly available; the PFC–CA1 cohort [28] (CRCNS *pfc-2*) is available from CRCNS on request to the original authors. Source data underlying every main-figure panel are provided with this paper.

## Code availability

NeuroGate is released at github.com/AliZareh-CoE/NeuroGate under the MIT license (release tag `v1.0.0`).

## Acknowledgements

We gratefully acknowledge the support of NVIDIA Corporation with the grant of the NVIDIA A100 GPU cloud compute credits used for this research. We thank the CRCNS, G-Node and OpenNeuro data archives, and the laboratories that openly shared the datasets analysed in this study.

## Author contributions

A.Z. conceived the study, designed and implemented the NeuroGate architecture, performed the experiments and analyses, and wrote the manuscript (Conceptualization; Methodology; Software; Formal analysis; Investigation; Data curation; Visualization; Writing — original draft). R.Y. contributed the neuroscience and biological design and concepts, and wrote and edited the manuscript (Conceptualization, biological and neuroscientific framing; Writing — original draft; Writing — review & editing). M.K.Ö. supervised the work and provided resources and project oversight as principal investigator (Supervision; Resources; Project administration; Writing — review & editing). All authors reviewed and approved the final manuscript.

## Ethics approval and consent to participate

This study is a secondary computational analysis of publicly available, previously published datasets; no new animal or human experiments were performed. All rodent and human recordings analysed here were acquired in the original studies under those studies’ institutional ethical approvals, and human data were collected with informed consent, as documented in the corresponding dataset publications [19, 28–32].

## References

[1] Kohn, A. et al. Principles of corticocortical communication: proposed schemes and design considerations. Trends Neurosci. 43, 725–737 (2020).

[2] Steinmetz, N. A., Zatka-Haas, P., Carandini, M. & Harris, K. D. Distributed coding of choice, action and engagement across the mouse brain. Nature 576, 266–273 (2019).

[3] Parvizi, J. & Kastner, S. Promises and limitations of human intracranial electroencephalography. Nat. Neurosci. 21, 474–483 (2018).

[4] Sip, V. et al. Data-driven method to infer the seizure propagation patterns in an epileptic brain from intracranial electroencephalography. PLOS Comput. Biol. 17, e1008689 (2021).

[5] Steinmetz, N. A. et al. Neuropixels 2.0: a miniaturized high-density probe for stable, long-term brain recordings. Science 372, eabf4588 (2021).

[6] Semedo, J. D., Zandvakili, A., Machens, C. K., Yu, B. M. & Kohn, A. Cortical areas interact through a communication subspace. Neuron 102, 249–259.e4 (2019).

[7] Semedo, J. D. et al. Feedforward and feedback interactions between visual cortical areas use different population activity patterns. Nat. Commun. 13, 1099 (2022).

[8] Pandarinath, C. et al. Inferring single-trial neural population dynamics using sequential auto-encoders. Nat. Methods 15, 805–815 (2018).

[9] Schneider, S., Lee, J. H. & Mathis, M. W. Learnable latent embeddings for joint behavioural and neural analysis. Nature 617, 360–368 (2023).

[10] Ye, J. & Pandarinath, C. Representation learning for neural population activity with Neural Data Transformers. Neurons Behav. Data Anal. Theory 5, 1–18 (2021).

[11] Jamison, K. W. et al. Krakencoder: a unified brain connectome translation and fusion tool. Nat. Methods 22, 1583–1592 (2025).

[12] Seth, A. K., Barrett, A. B. & Barnett, L. Granger causality analysis in neuroscience and neuroimaging. J. Neurosci. 35, 3293–3297 (2015).

[13] Friston, K. J., Harrison, L. & Penny, W. Dynamic causal modelling. NeuroImage 19, 1273–1302 (2003).

[14] Baccalá, L. A. & Sameshima, K. Partial directed coherence: a new concept in neural structure determination. Biol. Cybern. 84, 463–474 (2001).

[15] Luo, Z. et al. Mapping effective connectivity by virtually perturbing a surrogate brain. Nat. Methods 22, 1376–1385 (2025).

[16] d’Ascoli, S. et al. A foundation model of vision, audition, and language for in-silico neuroscience. Preprint at arXiv:2605.04326 (2026).

[17] Sweeney, S. T., Broadie, K., Keane, J., Niemann, H. & O’Kane, C. J. Targeted expression of tetanus toxin light chain in Drosophila specifically eliminates synaptic transmission and causes behavioral defects. Neuron 14, 341–351 (1995).

[18] Schiavo, G. et al. Tetanus and botulinum-B neurotoxins block neurotransmitter release by proteolytic cleavage of synaptobrevin. Nature 359, 832–835 (1992).

[19] Bolding, K. A. & Franks, K. M. Recurrent cortical circuits implement concentration-invariant odor coding. Science 361, eaat6904 (2018).

[20] von Ellenrieder, N., Beltrachini, L., Perucca, P. & Gotman, J. Size of cortical generators of epileptic interictal events and visibility on scalp EEG. NeuroImage 94, 47–54 (2014).

[21] Bai, S., Kolter, J. Z. & Koltun, V. An empirical evaluation of generic convolutional and recurrent networks for sequence modeling. Preprint at arXiv:1803.01271 (2018).

[22] Hochreiter, S. & Schmidhuber, J. Long short-term memory. Neural Comput. 9, 1735–1780 (1997).

[23] Dosovitskiy, A. et al. An image is worth 16 × 16 words: Transformers for image recognition at scale. Preprint at arXiv:2010.11929 (2020).

[24] van den Oord, A. et al. WaveNet: A generative model for raw audio. Preprint at arXiv:1609.03499 (2016).

[25] Schirrmeister, R. T. et al. Deep learning with convolutional neural networks for EEG decoding and visualization. Hum. Brain Mapp. 38, 5391–5420 (2017).

[26] Lawhern, V. J. et al. EEGNet: a compact convolutional neural network for EEG-based brain–computer interfaces. J. Neural Eng. 15, 056013 (2018).

[27] Izenman, A. J. Reduced-rank regression for the multivariate linear model. J. Multivariate Anal. 5, 248–264 (1975).

[28] Fujisawa, S., Amarasingham, A., Harrison, M. T. & Buzsáki, G. Behavior-dependent short-term assembly dynamics in the medial prefrontal cortex. Nat. Neurosci. 11, 823–833 (2008).

[29] Boran, E. et al. Dataset of human medial temporal lobe neurons, scalp and intracranial EEG during a verbal working memory task. Sci. Data 7, 30 (2020).

[30] Miller, K. J. A library of human electrocorticographic data and analyses. Nat. Hum. Behav. 3, 1225–1235 (2019).

[31] Delorme, A. EEG data from an auditory oddball task. OpenNeuro ds003061 (2020).

[32] Ajra, Z. et al. Multimodal dataset from the CMx7-MM experiment. OpenNeuro ds007554 v1.0.0 (2026).

[33] Owen, L. L. W. et al. A Gaussian process model of human electrocorticographic data. Cereb. Cortex 30, 5333–5345 (2020).

[34] Ronneberger, O., Fischer, P. & Brox, T. U-Net: Convolutional networks for biomedical image segmentation. Med. Image Comput. Comput.-Assist. Interv. (MICCAI) 234–241 (2015).

[35] Yu, F. & Koltun, V. Multi-scale context aggregation by dilated convolutions. Int. Conf. Learn. Represent. (ICLR) (2016).

[36] Hu, J., Shen, L. & Sun, G. Squeeze-and-excitation networks. Proc. IEEE Conf. Comput. Vis. Pattern Recognit. (CVPR) 7132–7141 (2018).

[37] Perez, E., Strub, F., de Vries, H., Dumoulin, V. & Courville, A. FiLM: Visual reasoning with a general conditioning layer. Proc. AAAI Conf. Artif. Intell. 32 (2018).

[38] Shapley, L. S. A value for n-person games. Contributions to the Theory of Games II (Princeton Univ. Press) 307–317 (1953).

[39] Kraskov, A., Stögbauer, H. & Grassberger, P. Estimating mutual information. Phys. Rev. E 69, 066138 (2004).

[40] Benjamini, Y. & Hochberg, Y. Controlling the false discovery rate: A practical and powerful approach to multiple testing. J. R. Stat. Soc. B 57, 289–300 (1995).

[41] Efron, B. & Tibshirani, R. J. An introduction to the bootstrap. Chapman & Hall/CRC, New York (1993).

