## Supplementary Tables S1-S8 for "NeuroGate: waveform translation between brain regions"

This Supplementary Information contains Supplementary Tables S1–S8: the exact values and exact $P$-values behind every quantitative claim in the main text. All $P$-values are exact rather than thresholded. Every value re-derives from the per-figure Source Data files that accompany the paper (`Source Data Fig. 1`, `Fig. 2` and `Fig. 3`, one Excel workbook per figure), and matches the main text.

There are no Supplementary Figures.

### Contents


**S1 — Per-pathway reconstruction statistics** **1**

**S2 — Baseline comparison** **2**

**S3 — Leave-one-component-out ablation** **2**

**S4 — Depth × width sweep** **3**

**S5 — TeLC perturbation experiments** **3**

**S6 — Exp 7 label-permutation null** **4**

**S7 — Natural coupling baseline** **4**

**S8 — Multi-source decomposition** **4**

**Source data** **6**


### Supplementary Table S1 — Per-pathway reconstruction statistics

All 112 pathway × direction × split cells (56 directed pathways, Test and Val). $n$ is the number of subjects or sessions contributing to that pathway; $P$ is a one-sided paired Wilcoxon signed-rank test of model $r$ against the natural inter-region coupling baseline on the same subjects, channels and windows. Anchors Fig. 2d–f.

The complete table — all 112 pathway × direction × split cells with per-pathway $n$, median model $r$, median natural-coupling $r$ and exact one-sided paired Wilcoxon $P$ — exceeds the one-page limit for a typeset table and is supplied as the machine-readable file `Supplementary_Table_S1_per_pathway.csv`.

**Table S1:** Per-pathway reconstruction statistics, summarised by dataset and split.

| Dataset group | Split | pathways | median model $r$ | median natural $r$ | exceeding baseline |
|---|---|---|---|---|---|
| rodent/sEEG/ECoG | test | 16 | 0.431 | 0.491 | 6/16 |
| rodent/sEEG/ECoG | val | 16 | 0.657 | 0.484 | 16/16 |
| P300-EEG | test | 20 | 0.518 | 0.442 | 19/20 |
| P300-EEG | val | 20 | 0.519 | 0.304 | 20/20 |
| CMx7-EEG | test | 20 | 0.604 | 0.478 | 20/20 |
| CMx7-EEG | val | 20 | 0.637 | 0.570 | 19/20 |

### Supplementary Table S2 — Baseline comparison

Held-out $R^2$ per leave-one-session-out fold on the rodent OB→PCx pathway (CRCNS *pcx-1*, $n = 9$ folds) for NeuroGate, nine deep-learning baselines and four linear baselines. CV is the across-fold coefficient of variation (SD / mean). NeuroGate's CV of 0.218 is the lowest of the ten architectures; every deep-learning baseline lies in 0.467–1.549. Anchors Fig. 1c.

**Table S2:** Baseline comparison: mean held-out $R^2$, SD and CV across nine folds.

| Architecture | mean $R^2$ | SD | CV |
|---|---|---|---|
| NeuroGate | 0.5169 | 0.1127 | 0.2180 |
| deepconvnet | 0.1733 | 0.2684 | 1.5488 |
| eegnet | 0.2462 | 0.2542 | 1.0328 |
| linear | 0.3793 | 0.1911 | 0.5039 |
| lstm | 0.4047 | 0.2265 | 0.5597 |
| shallowconvnet | 0.2738 | 0.2443 | 0.8922 |
| simplecnn | 0.3912 | 0.1827 | 0.4671 |
| tcn | 0.4406 | 0.2152 | 0.4884 |
| vit | 0.3997 | 0.2434 | 0.6090 |
| wavenet | 0.3984 | 0.2372 | 0.5953 |
| linear_instant | 0.3793 | 0.0939 | 0.2476 |
| ridge_instant | 0.3788 | 0.0937 | 0.2473 |
| reduced_rank | 0.3801 | 0.0931 | 0.2448 |
| wiener | 0.3798 | 0.1186 | 0.3122 |

Per-fold values for all nine leave-one-session-out folds are supplied as `Supplementary_Table_S2_baselines.csv`.

### Supplementary Table S3 — Leave-one-component-out ablation

Each row removes one component from the full NeuroGate configuration (depth = 2, width = 256) and re-runs all nine folds. $P$ is a *two-sided* paired Wilcoxon signed-rank test against the full model across the nine folds (`scipy.stats.wilcoxon`, default `alternative='two-sided'`). For $n = 9$ folds all differing in the same direction the exact two-sided $P$ is $2/2^9 = 0.0039$. Anchors Fig. 1e.

**Sign convention.** $\Delta R^2$ is signed as a *removal effect* — the change in $R^2$ when the component is taken out — so a component that helps carries a negative value; the main text quotes the same magnitudes as positive contributions.

**Table S3:** Leave-one-component-out ablation.

| Variant | mean $R^2$ | $\Delta R^2$ vs full | Wilcoxon $P$ |
|---|---|---|---|
| full | 0.5231 | +0.0000 | — |
| no_adaptive | 0.5146 | -0.0086 | 0.2031 |
| no_film | 0.5125 | -0.0106 | 0.2031 |
| no_noise_aug | 0.5155 | -0.0076 | 0.3008 |
| no_se | 0.5191 | -0.0040 | 0.5703 |
| residual_none | 0.5116 | -0.0115 | 0.1289 |
| single_dilation | 0.4515 | -0.0716 | 0.0039 |

Per-fold values are supplied as `Supplementary_Table_S3_LOCO_ablation.csv`.

### Supplementary Table S4 — Depth × width sweep

Nine-cell architecture sweep, mean test $R^2$ per cell across the nine folds. The carried-forward configuration is `arch_d2_w256` ($R^2 = 0.5231$). Anchors Fig. 1d.

**Table S4:** Depth × width sweep.

| Configuration | $n$ sessions | mean test $R^2$ | SD |
|---|---|---|---|
| arch_d2_w128 | 9 | 0.5055 | 0.1989 |
| arch_d2_w256 | 9 | 0.5231 | 0.1852 |
| arch_d2_w64 | 9 | 0.5030 | 0.1764 |
| arch_d3_w128 | 9 | 0.4918 | 0.1581 |
| arch_d3_w256 | 9 | 0.5054 | 0.1738 |
| arch_d3_w64 | 9 | 0.4952 | 0.1516 |
| arch_d4_w128 | 9 | 0.4873 | 0.1462 |
| arch_d4_w256 | 9 | 0.5053 | 0.1806 |
| arch_d4_w64 | 9 | 0.4789 | 0.1482 |

### Supplementary Table S5 — TeLC perturbation experiments

All seven TeLC experiments. Exp 1–3 are the unilateral-injection cohort, simul-trained; Exp 4–5 the Thy1 cohort trained between subjects; Exp 6–7 the Thy1 cohort with per-fold leave-one-subject-out fine-tuning. Exp 7 is the headline causal experiment. Anchors Fig. 3g–i.

†Exp 4 and Exp 5 are between-subjects comparisons and use a one-sided Mann–Whitney $U$ test (`scipy.stats.mannwhitneyu`, `alternative= 'greater'`) with Cohen's $d$ on the pooled SD, per Methods, *Statistics*; the remaining rows are paired tests with Cohen's $d_z$.

Exp 7 additionally carries a percentile bootstrap 95% CI of $[1.22, 3.02]$ (10,000 resamples, seed 42) and the label-permutation control tabulated in Supplementary Table S6.

**Table S5:** TeLC perturbation experiments, Exp 1–7.

| **Experiment** | **n** | **paired-t P** | **Wilcoxon P** | **Cohen's** $d\_z$ |
|---|---|---|---|---|
| Exp1 OB→PCx (unilateral-injection, simul) | 4 | 0.0205 | 0.0625 | 1.725 |
| Exp2 PCx→OB (unilateral-injection, simul) | 6 | 0.8843 | 0.8906 | -0.556 |
| Exp3 Y-junction (unilateral-injection) | 4 | 0.0163 | 0.0625 | 1.885 |
| Exp6 OB→PCx (Thy1 LOSO fine-tune) | 8 | — | 0.9258 | -0.531 |
| Exp7 PCx→OB (Thy1 LOSO fine-tune) | 8 | — | 0.0039 | 1.616 |
| Exp4 OB→PCx (Thy1, between-subjects) | 8 vs 15 | — | $0.9156^{\dagger}$ | -0.739 |
| Exp5 PCx→OB (Thy1, between-subjects) | 8 vs 15 | — | $0.9988^{\dagger}$ | -1.424 |

### Supplementary Table S6 — Exp 7 label-permutation null

The label-permutation control for the headline causal experiment (Exp 7, PCx→OB under TeLC silencing; master seed 42). $N = 100$ random Ctrl/TeLC label assignments were pushed through the identical per-fold fine-tuning pipeline; $k$ is the direction-consistency count, the number of leave-one-subject-out folds in which the silenced prediction degraded relative to control. Anchors Fig. 3i and Results §4.

**Table S6:** Exp 7 label-permutation null distribution.

| $k$ **(direction-consistency)** | **Permutations** ($N = 100$) | **Fraction** |
|---|---|---|
| 0 | 6 | 0.060 |
| 1 | 11 | 0.110 |
| 2 | 15 | 0.150 |
| 3 | 21 | 0.210 |
| 4 | 23 | 0.230 |
| 5 | 14 | 0.140 |
| 6 | 8 | 0.080 |
| 7 | 2 | 0.020 |
| 8 | 0 | 0.000 |

Observed $k = 8/8$; null mean $k = 3.28$, range 0–7. No permutation reached $k = 8$. Permutation $P = (\#\{k_{\text{perm}} \geq k_{\text{obs}}\} + 1)/(N + 1) = 0.0099$ (Phipson & Smyth 2010). Per-permutation $k$ values are in Source Data Fig. 3, sheet `Fig3_shuffle_k`.

### Supplementary Table S7 — Natural OB↔PCx coupling baseline

Model-free reference: real-vs-real Pearson $r$ and $R^2$ between the recorded olfactory bulb and piriform cortex signals, per session, on the same windows used to score the model. The MEAN row is the natural-coupling baseline reported in the main text ($R^2 = 0.115$). Anchors Fig. 1c (dotted line).

### Supplementary Table S8 — Multi-source decomposition

Per-target saturation-fit parameters for the two scalp cohorts. $p$ is the per-source independence parameter of the geometric fit $R^2(k) = R^2_\infty[1 - (1 - p)^k]$, a four-point two-parameter fit; $a(Y) = 1 - R^2_{\text{full}}$ is the intrinsic share. Anchors Fig. 3a–f.

**Table S7:** Natural OB↔PCx coupling baseline.

| **Session** | **natural coupling** $r$ | **natural coupling** $R^2$ |
|---|---|---|
| 160819 | 0.6145 | 0.2291 |
| 160820 | 0.7188 | 0.4377 |
| 170608 | 0.5379 | 0.0757 |
| 170609 | 0.5021 | 0.0042 |
| 170614 | 0.579 | 0.1579 |
| 170618 | 0.6446 | 0.2893 |
| 170619 | 0.502 | 0.004 |
| 170621 | 0.4615 | -0.077 |
| 170622 | 0.4556 | -0.0888 |
| MEAN (= manuscript natural-coupling baseline) | 0.5573 | 0.1147 |

**Table S8:** Multi-source decomposition per target.

| **Dataset** | **Target** | $n$ | **mean** $p$ | **mean** $R^2$__$\infty$ | **mean** $a(Y)$ | **median** $a(Y)$ |
|---|---|---|---|---|---|---|
| P300-EEG | frontal | 13 | 0.534 | 0.56 | 0.474 | 0.451 |
| P300-EEG | central | 13 | 0.566 | 0.555 | 0.46 | 0.452 |
| P300-EEG | parietal | 13 | 0.583 | 0.344 | 0.662 | 0.651 |
| P300-EEG | occipital | 13 | 0.549 | 0.312 | 0.696 | 0.722 |
| P300-EEG | temporal | 13 | 0.725 | 0.459 | 0.535 | 0.543 |
| CMx7-EEG | frontal | 12 | 0.546 | 0.742 | 0.292 | 0.329 |
| CMx7-EEG | central | 12 | 0.645 | 0.713 | 0.298 | 0.306 |
| CMx7-EEG | parietal | 12 | 0.665 | 0.553 | 0.448 | 0.477 |
| CMx7-EEG | occipital | 12 | 0.589 | 0.468 | 0.539 | 0.527 |
| CMx7-EEG | temporal | 12 | 0.726 | 0.553 | 0.443 | 0.436 |

The full per-subject $R^2(k)$ ladders, Shapley atoms and $5 \times 5$ unique-contribution matrices are in Source Data Fig. 3.

### Source data

Per-figure Source Data accompany the paper as one Excel workbook per main figure, each sheet labelled with the panel it supports:

- `Source Data Fig. 1` — 5 sheets: baseline comparison (S2), natural-coupling baseline (S7), depth × width sweep (S4), leave-one-component-out ablation (S3), and NeuroGate per-fold values across all datasets.
- `Source Data Fig. 2` — 10 sheets: per-pathway statistics (S1), the directional-network edge weights, the plotted exemplar waveforms, the 33-metric fidelity heatmap, per-fold phase–amplitude coupling, and the per-subject $r$ values for each raincloud panel.
- `Source Data Fig. 3` — 15 sheets: the multi-source decomposition (S8), per-subject $R^2(k)$ ladders and Shapley atoms, the $5 \times 5$ region matrices, the seven TeLC experiments (S5), the Exp 7 permutation null (S6), and the raw per-fold Ctrl/TeLC values for every panel.

The NeuroGate model, its training and evaluation code, and the analysis code for the experiments reported here are released at `github.com/AliZareh-CoE/NeuroGate` under the MIT licence (see Methods, *Code availability*).
